# A Systematic Multi-LLM AI Framework for Immunotherapy Biomarker Discovery and Target Identification

**DOI:** 10.1101/2025.06.10.658988

**Authors:** Namu Park, Jung Hyun Lee

## Abstract

Immune checkpoint inhibitors have revolutionized cancer therapy, yet their efficacy is frequently limited by primary and acquired resistance. This underscores the need for novel immune regulatory targets that can improve or broaden patient responsiveness to immunotherapy. Among these, immunoglobulin (Ig) domain-containing proteins play central roles in immune signaling and tumor immune evasion, but many remain underexplored as therapeutic targets. Although recent advances in RNA sequencing, single-cell technologies, and multi-omics have expanded our understanding of gene and protein expression, integrating these data into actionable therapeutic insights remains a major challenge.

Recent developments in Artificial Intelligence (AI), particularly in natural language processing and large language models (LLMs), offer a promising solution. Models such as GPT-4o, Llama 3.1–8B, and Gemini 1.5 Flash have demonstrated exceptional reasoning capabilities in biomedical applications. However, their use in immunotherapy biomarker discovery has yet to be fully realized.

In this study, we introduce a novel AI-based framework that employs a multi-LLM approach to systematically identify and prioritize candidate biomarkers from a curated set of Ig-domain-containing genes. By integrating multi-omics data with structured prompt engineering and comparative model reasoning, our platform enhances the robustness, reproducibility, and interpretability of gene selection while minimizing bias inherent to single-model predictions. To our knowledge, this is the first study to apply a multi-LLM strategy to immunotherapy biomarker discovery. Our findings support the broader utility of LLMs in precision oncology and highlight their potential to accelerate the identification of novel, clinically actionable targets.

## MAIN TEXT

Immune checkpoint inhibitors have revolutionized cancer therapy, yet their efficacy is frequently limited by primary and acquired resistance. This underscores the urgent need to discover novel immune regulatory targets that can improve or broaden patient responsiveness to immunotherapy. Biomarker discovery plays a pivotal role in this endeavor, enabling more precise patient stratification, prediction of therapeutic outcomes, and identification of resistance mechanisms. However, despite its clinical importance, effective strategies for systematically identifying robust and translational biomarkers remain underdeveloped. Immunoglobulin (Ig) domain-containing proteins, which play central roles in immune signaling and tumor immune evasion, remain underexplored despite their therapeutic potential. Although recent advances in RNA sequencing, single-cell analysis, and multi-omics technologies have expanded our knowledge of gene and protein expression, integrating these complex datasets into reproducible, clinically actionable biomarker discoveries continues to be a major challenge in the current biomedical landscape^1–3^.

Recent developments in Artificial Intelligence (AI), particularly in natural language processing and the advent of large language models (LLMs), present promising new opportunities for biomarker discovery. LLMs such as GPT-4o, Llama 3.1–8B, and Gemini 1.5 Flash have shown remarkable reasoning capabilities in biomedical contexts, sometimes even surpassing expert-level performance. Despite these advances, the application of LLMs in immunotherapy biomarker discovery has been limited, particularly with respect to systematic, multi-model approaches. In this study, we present a novel multi-LLM-based discovery framework that leverages the complementary strengths of different LLMs to prioritize candidate biomarkers from a curated list of Ig-domain-containing genes. By combining multi-omics data, structured prompt engineering, and comparative model reasoning, our approach enhances robustness, reproducibility, and interpretability while minimizing individual model biases. To our knowledge, this represents the first systematic application of a multi-LLM strategy to the field of immunotherapy biomarker discovery^4,5^.

For this analysis, we used publicly available bulk RNA-seq data from 84 prostate cancer patients (Van Espen et al., 2023), consisting of normalized expression values for 19,885 genes per sample. Focusing on immune-relevant targets, we selected 478 Ig-domain-containing genes, which include IG-like, V, I, C1, and C2 domain subtypes. Two gene subsets (each comprising the top 10% of genes ranked by average expression and expression variability, respectively) were generated, and overlapping genes were retained after removing HLA-related entries, resulting in a refined candidate list of 29 genes. This gene set served as the foundation for all subsequent analyses and model comparisons. Figure 1 illustrates the overall workflow of our platform, highlighting each step from data preprocessing and gene filtering to multi-LLM reasoning and final candidate prioritization. This gene set served as the foundation for all subsequent analyses and model comparisons.

**Figure 1.**
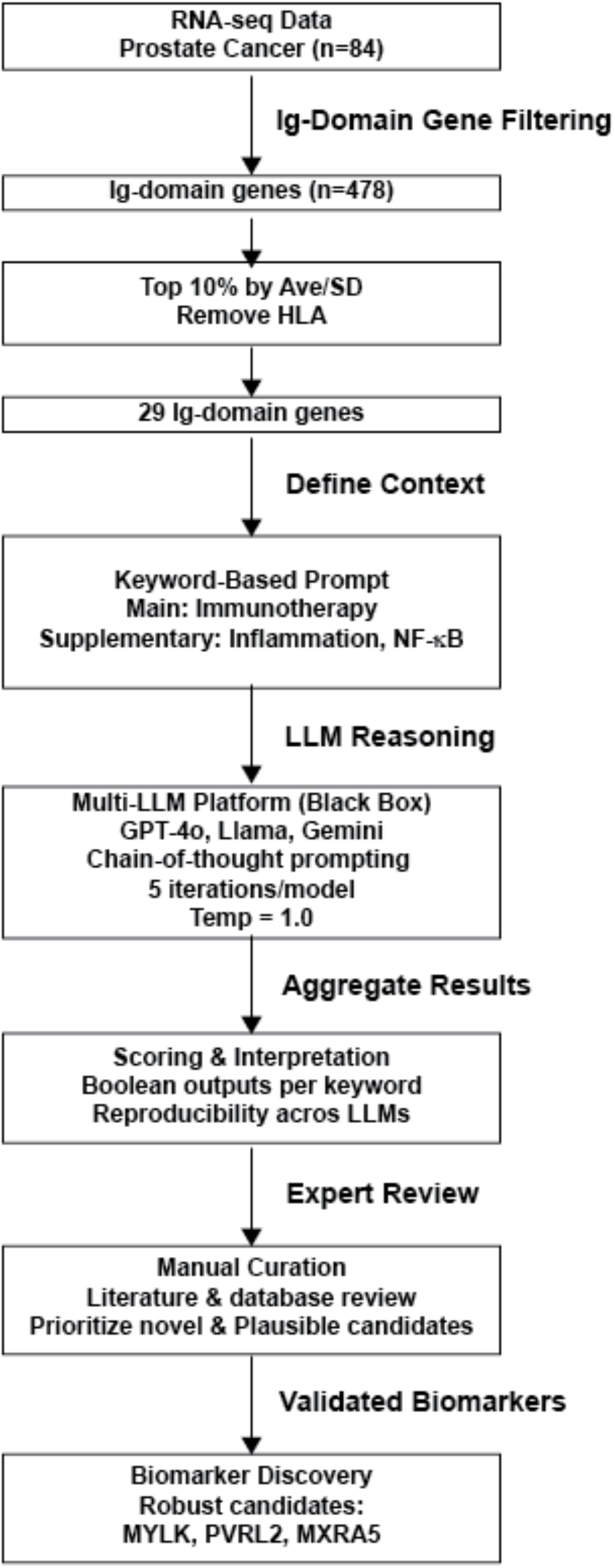
AI-driven framework for immunotherapy biomarker discovery using multi-LLM reasoning.This flowchart illustrates the sequential steps of our biomarker discovery pipeline. Starting from RNA-seq data derived from prostate cancer patients (n = 84), we filtered for immunoglobulin (Ig) domain-containing genes and selected top candidates based on expression metrics. A keyword-based prompt engineering strategy was applied, incorporating a main keyword (“immunotherapy”) and two supplementary keywords (“inflammation” and “NF-κB”), to structure the query inputs. These were processed through three independent Large Language Models (LLMs)GPT-4o, Llama 3.1–8B, and Gemini using consistent parameters and five inference iterations per model. The outputs were scored and classified according to their relevance to each keyword. Final candidate selection was refined through manual curation involving literature and database validation. This integrated platform enabled the identification of novel, biologically plausible immunotherapy biomarkers such as MYLK, PVRL2, and MXRA5, demonstrating the utility of multi-LLM reasoning in scalable, high-throughput biomarker discovery.

To ensure a systematic and interpretable prioritization of these genes, we designed a keyword-based prompt engineering method that incorporates three biologically meaningful dimensions: “immunotherapy,” “inflammation,” and “NF-κB signaling.” Each gene was evaluated independently by the three LLMs using identical input conditions. For each category, models assigned a relevance score from 1 (not related) to 3 (highly related), accompanied by concise rationales based on publicly available gene function descriptions. The prompts were structured to mimic expert-level review and were designed to be adaptable across domains and LLM architectures. This strategy minimizes the need for advanced technical expertise, allowing domain researchers to generate high-quality outputs using biologically grounded inputs. The full prompt structure and scoring system used in our evaluation are detailed in Supplementary Data 1.

Each of the 29 genes was evaluated using GPT-4o, Llama 3.1–8B, and Gemini 1.5 Flash under uniform conditions, including standardized input templates and five inference runs per model with a fixed temperature parameter of 1.0 to reduce stochastic variability. While GPT-4o and Gemini exhibited near-identical output in both scoring and rationale structure, Llama’s results, although directionally consistent, showed modest variation in score distribution. This divergence likely reflects differences in model architecture, training datasets, and domain-specific knowledge encoding. The overall consistency between GPT-4o and Gemini provides a strong internal validation for the multi-LLM design, while Llama’s variability underscores the potential for complementary insights.

Top-ranked genes identified by cross-model consensus included IL1R1, MYLK, PTPRS, MXRA5, and PVRL2. These genes were consistently prioritized across all models and scored highly in two or more categories. Importantly, these candidates emerged from a consensus-driven integration of three independent LLMs, demonstrating the strength of a multi-model approach to biomarker discovery. While genes like CD276 and B2M have well-established roles in immunotherapy, others such as MYLK, MXRA5, and PVRL2 have not yet been developed into immunotherapy drugs. Nevertheless, they exhibit strong mechanistic relevance to immune modulation and tumor progression. For example, PVRL2 has been associated with immune evasion and poor prognosis, while MXRA5 has been implicated in extracellular matrix remodeling and immune suppression. Furthermore, these genes were consistently upregulated across prostate cancer patient samples, adding weight to their biological relevance. Table 1 summarizes these prioritized targets, emphasizing that they are not only computationally validated by multiple LLMs but also represent novel, underutilized opportunities in immunotherapy development for prostate cancer. Table 2 presents the detailed scoring results derived from the structured prompt evaluations across the three LLMs, highlighting how these genes scored consistently high in immunotherapy-related categories and were upregulated in prostate cancer samples, underscoring their translational potential.

**Table 1.** LLM-Prioritized Gene Targets This table summarizes the top-ranked genes prioritized by multi-LLM consensus. Genes such as IL1R1, MYLK, and PTPRS show high agreement across all LLMs and have not been previously used in immunotherapy. Their potential biological functions suggest strong relevance for future therapeutic development.

| Gene | LLM Agreement | Prior Use in Immunotherapy | Potential Role |
| --- | --- | --- | --- |
| IL1R1 | High | No | Inflammatory cytokine receptor |
| MYLK | High | No | Endothelial permeability & immune infiltration |
| PTPRS | High | No | Tyrosine phosphatase with checkpoint potential |
| MXRA5 | High | No | ECM remodeling and immune suppression |
| PVRL2 | High | No | Immune evasion via T cell interaction |
| CD276 | High | Yes | Known checkpoint regulator (B7-H3) |
| B2M | High | Yes | MHC-I complex component |

**Table 2.** LLM-Based Gene Scoring Summary Across Three Immunological Dimensions This table presents average relevance scores for the top-ranked Ig-domain-containing genes, evaluated using three independent large language models (LLMs): GPT-4o, Llama 3.1, and Gemini 1.5 Flash. Each gene was assessed across three biologically meaningful categories—Immunotherapy (I), Inflammation (Inf), and NF-κB signaling (NFkB)—using structured, expert-level prompts. Scores range from 1 (not related) to 3 (highly related) and represent the mean of five inference runs per model with standardized input parameters. This comparative scoring framework enables robust and interpretable prioritization of candidate biomarkers for immunotherapy development.

| Gene | GPT4o [I] | GPT4o [Inf] | GPT4o [NFκB] | Llama3 [I] | Llama3 [Inf] | Llama3 [NFκB] | Gemini [I] | Gemini [Inf] | Gemini [NFκB] |
| --- | --- | --- | --- | --- | --- | --- | --- | --- | --- |
| B2M | 3 | 2 | 1.4 | 1.8 | 1.4 | 1.8 | 3 | 2 | 1 |
| CD276 | 3 | 2 | 1.2 | 2.4 | 2.2 | 1.8 | 2.4 | 2 | 1 |
| TAPBP | 3 | 1.2 | 1.2 | 1.4 | 1.2 | 1.2 | 3 | 2 | 1.2 |
| ALCAM | 2.8 | 2 | 1.2 | 1.6 | 2.6 | 1.6 | 2.8 | 2 | 1 |
| IL1R1 | 2.6 | 3 | 3 | 2.2 | 3 | 3 | 2 | 3 | 3 |
| PVRL2 | 2.6 | 2 | 1 | 2.4 | 1.2 | 1.6 | 3 | 2 | 1.2 |
| BSG | 2.2 | 3 | 1.8 | 1.6 | 2.2 | 1.4 | 2 | 2 | 1 |
| LRIG1 | 2 | 1.8 | 1 | 1.6 | 1.4 | 1 | 2.8 | 2 | 1 |
| PIGR | 2 | 2.2 | 1 | 2 | 1.4 | 1.2 | 2 | 2 | 1 |
| IGFBP7 | 1.8 | 2.6 | 1.2 | 1.4 | 2.2 | 1.2 | 2 | 2 | 1 |

Note: Genes were evaluated by GPT-4o, Llama3, and Gemini based on their relevance to Immunotherapy (I), Inflammation (Inf), and NF-κB signaling (NFκB) using a 3-point Likert scale. Scores represent the average from 5 inference runs per model.

To contextualize each candidate’s role in immune regulation, we conducted secondary analyses using established biological databases. Structural features were annotated using UniProt, InterPro, and the Protein Data Bank (PDB), with emphasis on conserved Ig-domain motifs and homology to known checkpoint molecules such as PD-1 and CTLA-4. Interaction networks were analyzed through STRING and BioGRID to identify intracellular signaling pathways linked to immune activation or suppression, particularly NF-κB, PI3K/AKT, and TCR signaling cascades. Expression data from GeneCards, Human Protein Atlas, and GTEx were used to evaluate each gene’s relevance across immune cell populations and tumor tissues, prioritizing those expressed on the cell surface and within immune-infiltrated tumor microenvironments.

We further propose a comprehensive validation pipeline to test the biological function of high-priority genes. First, RNA-seq data from TCGA and ICGC will be used to assess differential expression in immune-hot versus immune-cold tumors. In vitro experiments involving CRISPR-Cas9 knockout and cDNA overexpression in prostate cancer cell lines (e.g., PC3, DU145) will evaluate effects on immune signaling, including cytokine secretion and T cell activation. In vivo validation will involve gene-modified murine models such as RM-1 to assess tumor growth dynamics and response to anti-PD-1 therapy. Finally, integrative multi-omics profiling using ATAC-seq, proteomics, and single-cell RNA-seq will provide additional mechanistic insights into pathway modulation and tumor-immune interactions.

This multi-stage validation strategy ensures that candidate genes identified through LLM-based reasoning are biologically meaningful and clinically actionable. Our findings demonstrate that AI-generated hypotheses, when combined with biological data and expert interpretation, can support scalable and systematic biomarker discovery. The framework is generalizable beyond prostate cancer and can be applied to other tumor types or molecular families by modifying input parameters and prompt structures. Future iterations may incorporate feedback loops, domain-specific LLM tuning, and real-time integration with experimental platforms to further enhance predictive accuracy and translational relevance.

Ultimately, this work highlights the power of LLMs to serve as intelligent collaborators in the discovery of immunotherapy targets. By enabling high-throughput, interpretable prioritization of biologically relevant genes, our approach lays the groundwork for a new generation of AI-assisted tools in precision oncology.

## Supplementary Data 1

### Prompt Description

You are a biologist with expertise in immunotherapy, inflammation, and NF-κB signaling pathways. Using your expertise, analyze each gene from the input to assess their involvement in the following areas:

1. Immunotherapy
2. Inflammation
3. NF-κB signaling

Each gene should be evaluated using the provided function descriptions and relevant publications. Assign a relevance score for each category using a 3-point Likert scale:

‐ 3 = highly related
‐ 2 = partially related
‐ 1 = not related

In addition to numerical scores, provide a brief expert-level rationale (maximum 10 sentences) justifying the assigned scores. The output must follow the strict format below and should not include any other content:

***{‘GeneName’: [Immunotherapy_score, Inflammation_score, NFkB_score, “Rationale”]}***

### Example Evaluation

Gene: B2M

Function: Component of the class I major histocompatibility complex (MHC). Involved in the presentation of peptide antigens to the immune system. Exogenously applied M. tuberculosis EsxA or EsxA-EsxB (or EsxA expressed in host) binds B2M and decreases its export to the cell surface (total protein levels do not change), probably leading to defects in class I antigen presentation (PubMed:25356553). B2MG_HUMAN, P61769.

### Relevant Publications

‐ Craters on the melanoma surface facilitate tumor-immune interactions and demonstrate pathologic response to checkpoint blockade in humans. bioRxiv [Preprint]. 2024 Sep 19:2024.09.18.613595. doi: 10.1101/2024.09.18.613595
‐ An improved approach to generate IL-15+/+/TGFβR2−/− iPSC-derived natural killer cells using TALEN. Chen, A.P. et al. Cell Reports Methods. 2024 Sep; 4(9):100857.

### Output Format Example

{‘B2M’: [3, 2, 1, “B2M is a crucial component of MHC class I, directly involved in antigen presentation, a central process for immunotherapy. Its role in antigen presentation makes it highly relevant to immunotherapy (3). Altered B2M expression or function can impact immune responses and inflammation (2). However, there’s no direct evidence linking B2M to the NF-kB pathway itself (1). The cited publications on melanoma and NK cells highlight the importance of MHC class I in immune responses relevant to immunotherapy. Disruption of MHC class I antigen presentation can indirectly influence inflammatory processes. There is no direct mechanistic link between B2M and NF-kB activation or inhibition.”]}

